# Detection of mitotic neuroblasts provides additional evidence of steady state neurogenesis in the adult small intestinal myenteric plexus

**DOI:** 10.1101/2022.11.14.516462

**Authors:** Anastazja M. Gorecki, Jared Slosberg, Su Min Hong, Philippa Seika, Srinivas Puttapaka, Blake Migden, Anton Gulko, Alpana Singh, Chengxiu Zhang, Rohin Gurumurthy, Subhash Kulkarni

## Abstract

Maintenance of normal structure of the enteric nervous system (ENS), which regulates key gastrointestinal functions, requires robust homeostatic mechanisms, since by virtue of its location within the gut wall, the ENS is subject to constant mechanical, chemical, and biological stressors. Using transgenic and thymidine analogue-based experiments, we previously discovered that neuronal turnover – where continual neurogenesis offsets ongoing neuronal loss at steady state – represents one such mechanism. Although other studies confirmed that neuronal death continues into adulthood in the myenteric plexus of the enteric nervous system (ENS), the complicated nature of thymidine analogue presents challenges in substantiating the occurrence of adult neurogenesis. Therefore, it’s vital to employ alternative, well-recognized techniques to substantiate the existence of adult enteric neurogenesis in the healthy gut. Here, by using established methods of assessing nuclear DNA content and detecting known mitotic marker phosphor-histone H3 (pH3) in Hu^+^ adult ENS cells, we show that ∼10% of adult murine small intestinal myenteric Hu^+^ cells, and ∼20% of adult human small intestinal myenteric Hu^+^ cells show evidence of mitosis and hence are cycling neuroblasts. We observe that proportions of Hu^+^ cycling neuroblasts in the adult murine ENS neither vary with ganglia size, nor do they differ significantly between two intestinal regions – duodenum and ileum, or between sexes. Confocal microscopy provides further evidence of cytokinesis in Hu^+^ cells. The presence of a significant population of cycling neuroblasts in adult ENS provide further evidence of steady state neurogenesis in the adult ENS.

## Introduction

The cells of the enteric nervous system (ENS) are located entirely within the gut wall and are routinely exposed to significant and continual mechanical, chemical, and biological insults(1–3). Despite these ongoing and significant stressors, the question of how the adult ENS maintains itself has not been adequately addressed, especially given prior reports from multiple groups that a significant proportion of enteric neurons undergo apoptosis at steady state(4–6).

In a previous study, we reported the presence of a neurogenic mechanism, which gives rise to a significant number of new neurons at steady state in the adult gut(7). Using thymidine analogue studies and multiple lines of evidence to test for neuronal apoptosis, we demonstrated the existence of an adult enteric neurogenic program in the adult murine myenteric plexus that generates new neurons to replace dying apoptotic neurons, which account for ∼10% of all myenteric neurons at any given time. We proposed that this is one mechanism to maintain the structural and functional integrity of the adult ENS, especially in the myenteric plexus tissue. While other studies provide independent validation of the high rate of neuronal apoptosis in the adult ENS(6, 8–11), the complicated nature of thymidine-analogue based assays and their susceptibility to variations in tissue handling and fixation has made it difficult for investigators to consistently detect neurogenesis(12). Thus, there is a need to use alternate methods that can aid in the detection of adult neurogenesis and provide further evidence of a high rate of neurogenesis that counteracts the equally high rate of neuronal apoptosis.

In addition, we and others also recently showed the presence of significant heterogeneity in myenteric ganglia (13, 14). Myenteric ganglia exist in various sizes, as defined by their neuronal numbers, and in the small intestine can contain from 3 to 150 neurons (13). The relative distribution of ganglia size in the intestine was skewed significantly towards smaller ganglia (i.e. containing fewer neurons) (13). Similarly, proportions of various neuronal subpopulations contained within these ganglia show significant differences across regions of the intestine(14).

This heterogeneity begs the question of whether the rates of neurogenesis and neuronal loss are equally represented in the diverse ganglia of myenteric plexus in differing locations in the small intestine, and across the two sexes. Since thymidine analogue-based assays, which detect the presence of these chemicals dosed to animals over several days, do not permit the study of ongoing neurogenesis at a defined snapshot of time, these assays are unsuitable for providing a real time assessment of putative differences in neurogenic niches across ganglia and locations. Further, despite high rates of apoptosis (which range from 5 – 30% of all myenteric neurons) reported in human ENS (4, 5, 9), the presence of a neurogenic mechanism in the human ENS is unclear. Since thymidine analogues cannot be administered to human beings, these assays cannot be translated to human tissues, and thus, there is a need to simplify the protocols to facilitate better assessment of neurogenesis both in mice and in human specimens.

To address these issues, in this report, we use three different methods to interrogate whether neurogenesis occurs in adult healthy small intestinal ENS. First, using high-resolution confocal microscopy, we find evidence of bi-nucleated Hu-immunolabeled ganglionic cells in the adult small intestinal myenteric plexus. Second, by performing a flow cytometry-based estimation of DNA content in myenteric Hu-immunolabeled nuclei, we observe DNA content suggestive of S-phase and G2/M phase in significant proportions of Hu-immunolabeled nuclei – which are indicative of active DNA synthesis and mitosis in a population of adult murine myenteric neurons. Finally, by performing immunostaining with an established mitotic marker phosphor-histone H3 (pH3), we identified the presence of mitotic Hu-immunolabeled cells in the adult murine and human ENS. The proportions of pH3-immunoreactive neurons in the adult murine myenteric plexus were found to match the previously observed proportions for apoptotic neurons and were conserved between myenteric ganglia regardless of their size, location in the small intestine, and the sex of the animals. Thus, these three different methods together provide additional evidence of continuous neurogenesis in the adult healthy gut.

## Methods

### Animals

Experimental protocols were approved by The Johns Hopkins University’s Animal Care and Use Committee and the Animal Care and Use Committee at Beth Israel Deaconess Medical Center, in accordance with the guidelines provided by the National Institutes of Health. Age matched 2 – 3-month-old adult wildtype C57BL/6 mice were used.

### Isolation and immunolabeling of neuronal nuclei from adult longitudinal muscle – myenteric plexus (LM-MP) tissue

Mice were anesthetized with isoflurane and euthanized by cervical dislocation. For small intestinal tissue isolation, a laparotomy was performed, and the entire small intestine was removed and lavaged with PBS containing penicillin-streptomycin (Pen-Strep; Invitrogen). The small intestine was then cut into 1-cm-long segments. Next, tissues were placed over a sterile plastic rod and a superficial longitudinal incision was made along the serosal surface and the LM-MP was peeled off from the underlying tissue using a wet sterile cotton swab (7) and directly frozen in liquid nitrogen and stored at −80^ο^C for further processing. Two different protocols were used hence-forth, which are detailed as follows.

BIDMC protocol: Frozen tissue was next transferred to a gentleMACs C-Tube (Miltenyi Inc.) containing with 2 ml of TST buffer (containing 60 μl of 5M NaCl, 42 μl of 1M MgCl_2_, 20 μl of 1M Tris HCl pH 7.5, 2 μl of 1M CaCl_2_, 10 μl 2% Bovine Serum Albumin (BSA), 6 μl 10% Tween-20, in 1860 μl ultrapure water) and allowed to thaw for 2 minutes. Thawed tissues were chopped using dissecting scissors for 45 seconds, and then added to the C-tube, which is installed on the gentleMACs dissociator (Miltenyi Inc.). The tissue was processed through two rounds of 268 rotations, for a total of 72 seconds dissociation time after which the tube was placed and incubated on ice for 10 min. Next, using a wide-bore, low retention 1000 μl pipette tip (Olympus, Cat. No. 23-165RL), the suspension was transferred to a 40 μm pre-wet cell sieve (Fisher Scientific Cat. No.22-363-547) that was set on top of a sterile 50 ml falcon tube and filtered through. The C-tube was washed with 1 ml of 1X ST buffer (TST buffer minus BSA and Tween-20), and the buffer was next transferred to the cell sieve. The cell sieve was then washed with 2 ml of 1X ST buffer and then discarded. Using again the wide-bore low retention 1000 μl pipette tip (Olympus, Cat. No. 23-165RL), the filtered nuclei suspension was transferred to a pre-wet 40 μm cell sieve (Fisher Scientific Cat. No.22-363-547) that is set on top of a 5 ml FACS tube. The suspension was spun down in a swing bucket centrifuge at 500g for 5 minutes and at 4^ο^C. Centrifuge brake was set at half to ensure that the pellet is not disturbed. The top ∼4.9 ml of the supernatant was removed, and the pellet was washed with 5 ml PBS solution containing 0.02% BSA, which was added to the tube gently and the tube was again spun at 500g for 5 minutes and at 4^ο^C. The supernatant was removed, and the nuclear pellet was resuspended in 500 μl of PBS containing 0.02% BSA. The suspension was then split equally into two tubes. The directly conjugated antibody Anti-Hu 488 (1:750, Abcam# ab237234) was added to the first tube, while the second tube served as the control. A drop of NucBlue (Thermofisher) was added to both the tubes. Both the tubes were incubated on ice for 45 minutes, after which the nuclear suspensions were spun down at 500g for 5 minutes and at 4^ο^C, with the centrifuge brakes again set to half. 450 μl of supernatant from each tube was removed and the nuclear pellet was resuspended in 450 μl of PBS solution containing 0.02% BSA. The suspensions were then analyzed for NucBlue fluorescence intensity (as a marker for DNA content) in Hu-immunofluorescent nuclei using the CytoFLEX flow cytometer.

JHU protocol: Frozen and chopped tissue was placed into an ice-cold gentleMACS C-tube with 3 ml of TST buffer. Tissue was dissociated on a gentleMACS Octo Dissociator using the *4C_nuclei_01* protocol with pre-chilled OctoCooler. Disrupted cell suspension was subsequently incubated on ice for 5 min before being filtered through a 40 μm pre-wet (with 1X ST buffer) cell sieve that is set on top of a 50 ml falcon tube. The C-tube was rinsed with 1 ml ST buffer and an additional 2 ml of 1X ST buffer was used to rinse the filter. The nuclei suspension was centrifuged in a swing bucket centrifuge at 500g for 5 minutes and at 4^ο^C with the centrifuge brakes set to half. The pellet was resuspended in 5 ml PBS containing 1% BSA, centrifuged and pelleted again, and finally resuspended in PBS containing 1% BSA and 5% normal goat serum (NGS). This suspension was incubated on ice for 5 minutes after which the suspension is divided equally into two tubes. Anti-Hu 647 directly conjugated antibody (Abcam, ab237235; RRID: AB_3668731) was added at a concentration of 1:250 to the first but not the second tube and both the tubes are incubated on ice for 45 minutes. Excess antibody was removed by washing the suspension twice (again with centrifugation at 500g for 5 minutes and at 4^ο^C with the centrifuge brakes set to half) and the nuclei suspensions were finally suspended in 1 ml sterile PBS in a 15 ml conical tube. Next, a fixative was freshly prepared by adding 50 μl of 50 mg/ml dithiobis(succinimidyl propionate) (DSP) in dimethyl sulfoxide (DMSO) solution to 4 ml of ice-cold methanol, dropwise while vortexing at moderate speed. The nuclei suspensions were fixed by adding the freshly prepared fixative dropwise to the nuclei, while swirling the suspension to reduce clumping. The fixed nuclei were next incubated at 4^ο^C for 10 minutes on an end-over-end rotor (speed of 15 rpm). Fixed nuclei were rehydrated by adding 2 volumes of PBS containing 0.1% Triton-X, and then washed twice with PBS containing 1% BSA. DAPI was added to both tubes at a concentration of 1 μg/ml, and samples were filtered through a 40 μm before analyses on a flow analyzer (BD FACSAria III).

Flow data was analyzed on Flowjo 10.

### Tissue preparation

Mice were anesthetized with isoflurane and sacrificed by cervical dislocation. For small intestinal tissue isolation, a laparotomy was performed, and the entire small intestine was removed and lavaged with PBS containing penicillin-streptomycin (Pen-Strep; Invitrogen), then cut into 2-cm-long segments, such that segments from duodenum, jejunum, and ileum were separated. Next, tissues were placed over a sterile plastic rod and a superficial longitudinal incision was made along the serosal surface and the longitudinal muscle containing myenteric plexus (LM-MP) was peeled off from the underlying tissue using a wet sterile cotton swab (7) and placed in Opti-MEM medium (Invitrogen) containing Pen-Strep. The tissue was then laid flat and fixed with freshly prepared ice cold 4% paraformaldehyde (PFA) solution for 4-5 minutes in the dark. For a small subset of tissues, where we tested how fixation alters immunoreactivity, we fixed LM-MP tissues for 10, 15, and 20 minutes with freshly prepared ice cold PFA solution. All LM-MP tissues were fixed within 30 minutes of sacrifice. After the fixation, the tissue was stored in ice cold sterile PBS with Pen-Strep for immunofluorescence staining and subsequent microscopy.

### Immunochemistry

For murine tissue: The fixed LM-MP tissue was washed twice in ice-cold PBS in the dark at 16°C. The tissue was incubated in blocking-permeabilizing buffer (BPB; 5% normal goat serum with 0.3% Triton-X) for 1 hour, followed by incubation with either the combination of (a) anti-pH3 antibody (Unconjugated antibody EMD Milipore # 06-570; RRID:AB_310177, 1:500; Conjugated antibody, #3458S, Cell Signaling; 1:250; RRID:AB_10694086) and ANNA-1 (1:1000; patient antisera against the neuronal Hu antigens; RRID:AB_2813895), (b) anti-Cleaved Caspase 3 (#Asp175, Cell Signaling, 1:250; RRID:AB_2070042) and ANNA-1(1:1000), (c) anti-PGP9.5 (#ab108986, Abcam, 1:250, RRID:AB_10891773) and ANNA-1(1:1000), (d) anti-Hu (#ab184267, Abcam, 1:1000, RRID:AB_2864321) and ANNA-1(1:1000), and (e) anti-Nestin (#NB100-1604, Novus; 1:250; RRID:AB_2282642) and anti-Hu (#ab184267, Abcam, 1:1000) for 48 h at 16°C in the dark with shaking at 55 rpm. The tissues were then washed three times (15-min wash each) in PBS at room temperature in the dark and incubated in the appropriate secondary antibody (Anti-human 488; RRID:AB_2536548, Anti-Chicken 488; RRID:AB_142924, and Anti-rabbit 647; RRID:AB_2535813, Invitrogen, all at 1:500) at room temperature for 1 hour while on a rotary shaker (65 rpm). The tissues were again washed three times in PBS at room temperature, counterstained with DAPI to stain the nuclei, overlaid with Prolong Antifade Gold mounting medium, cover-slipped, and imaged. ANNA1 patient antiserum was a kind gift from Dr. Sean Pittock at Mayo Clinic.

For formalin fixed paraffin embedded (FFPE) human tissue: Use of commercially procured human tissue was approved by the Institutional Review Board of the Beth Israel Deaconess Medical Center. FFPE tissue sections from human full thickness duodenal tissue of 4 different patients (2 Males and 2 Females) with no known GI dysmotility disorder were procured from a commercial vendor and from pathology archive at BIDMC. To process them, slides were first baked at 55^ο^C for 15 minutes, after which, they were deparaffinized and rehydrated by immersion through the following solutions: 1) Two washes (5 minutes each) with Xylene; 2) two washes with 100% ethanol (5 minutes each); 3) two washes with 95% ethanol (5 minutes each); 4) two washes with 70% ethanol (5 minutes each); and finally 6) two washes with deionized water (5 minutes each). Next, using Sodium Citrate buffer (pH 6.0), 40-minute antigen retrieval was performed in a pressure cooker. The slides were then cooled in ice-cold 1X PBS. Slides were marked with a hydrophobic pen around the sections, and then blocked and permeabilized in BPB for 1 hour. Buffer was washed off with 1X PBS and sections were incubated with either the combination of directly conjugated anti-pH3 antibody (EMD Milipore, 1:250) and directly conjugated anti-Hu antibody (1:300), or the combination of anti-Cleaved Caspase 3 (CC3; 1:250) and ANNA-1 (1:500) for 24 h at 16°C in the dark. For tissue sections treated with anti-CC3 and anti-ANNA1 antibodies, following incubation with primary antibody, the sections were washed three times for 15-minutes each in PBS at room temperature in the dark, then incubated with the appropriate secondary antibodies (Invitrogen Anti-human 488 and Anti-rabbit 647; both at 1:500) at room temperature for 1 hour. Next, the sections were again washed three times in PBS at room temperature, counterstained with DAPI to stain the nuclei, overlaid with Prolong Antifade Gold mounting medium, cover-slipped, and imaged.

### Microscopy

Microscopy was performed using the oil immersion 40X objective on the Olympus Fluoview 3000rs confocal microscope with galvano and resonance scanning mode, with 40X water immersion objective on the Zeiss LSM880, and with 20X and 40X magnification of the EVOS M7000 microscope. Control tissues that were immunostained with secondary antibodies only were used to set the threshold laser intensities. In mice, all ANNA-1/Hu-immunostained neurons were included in the analysis. Image analyses was performed by using Fiji (NIH) or ImarisViewer (10.2.0).

## Results

### Hu-expression marks a subset of multi-nucleated ganglionic cells in adult small intestinal murine myenteric ganglia

Hu-expression is known to be a pan-neuronal marker for all adult enteric neurons (13, 15–19). These studies have used both commercially available antibodies against the neuronal Hu antigens (HuB/C/D), as well as the patient-derived ANNA1 sera that contain autoantibodies against the neuronal Hu antigens. As these are often used as alternatives, we tested whether they detected the same ganglionic cells and found by co-immunostaining that the commercially available anti-Hu antibodies detect the same cells as the anti-Hu autoantibodies in the patient-derived ANNA1 antisera (Fig 1A). Next, we tested the co-expression of Hu and the pan-neuronal marker PGP9.5 and found that Hu immunostaining (ANNA1) co-localized with PGP9.5 immunostaining in the adult murine small intestinal myenteric plexus (Fig 1B). By imaging 3-dimensional (3D) z-stack of a Hu and PGP9.5-immunostained adult murine small intestinal myenteric ganglia, we found that that some Hu^+^ PGP9.5^+^ cells within myenteric ganglia contained more than 1 nucleus (Fig 2). This observation was reproduced in tissues from another mouse where two nuclei were observed in a single Hu-expressing cell (Suppl. Fig 1).

**Figure 1:**
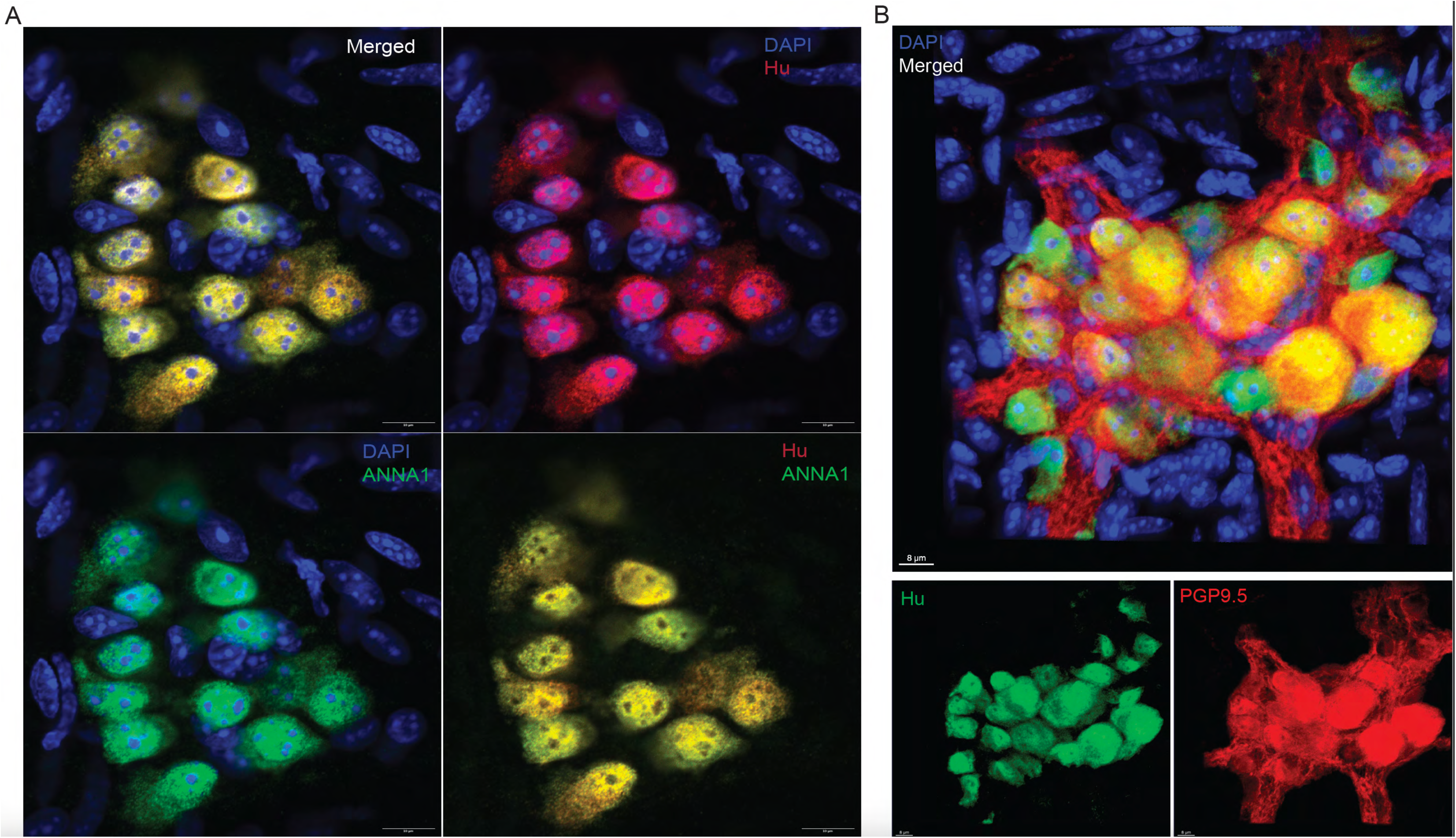
Hu-expression labels all myenteric ganglia cells that express the pan-neuronal marker PGP9.5. (A) Representative image (merged and color-segregated) showing co-immunolabeling of an adult murine small intestinal myenteric ganglion with commercially available anti-Hu antibody (red) along with anti-Hu antibodies in the patient-derived ANNA1 antisera (green). Nuclei are labeled with DAPI (blue). Scale bar denotes 10 μm. (B) Representative image (merged and panel-segregated) showing co-immunolabeling of an adult murine small intestinal myenteric ganglion with commercially available antibodies against Hu (green) and PGP9.5 (red). Nuclei are labeled with DAPI (blue). Scale bar denotes 8 μm.

**Figure 2:**
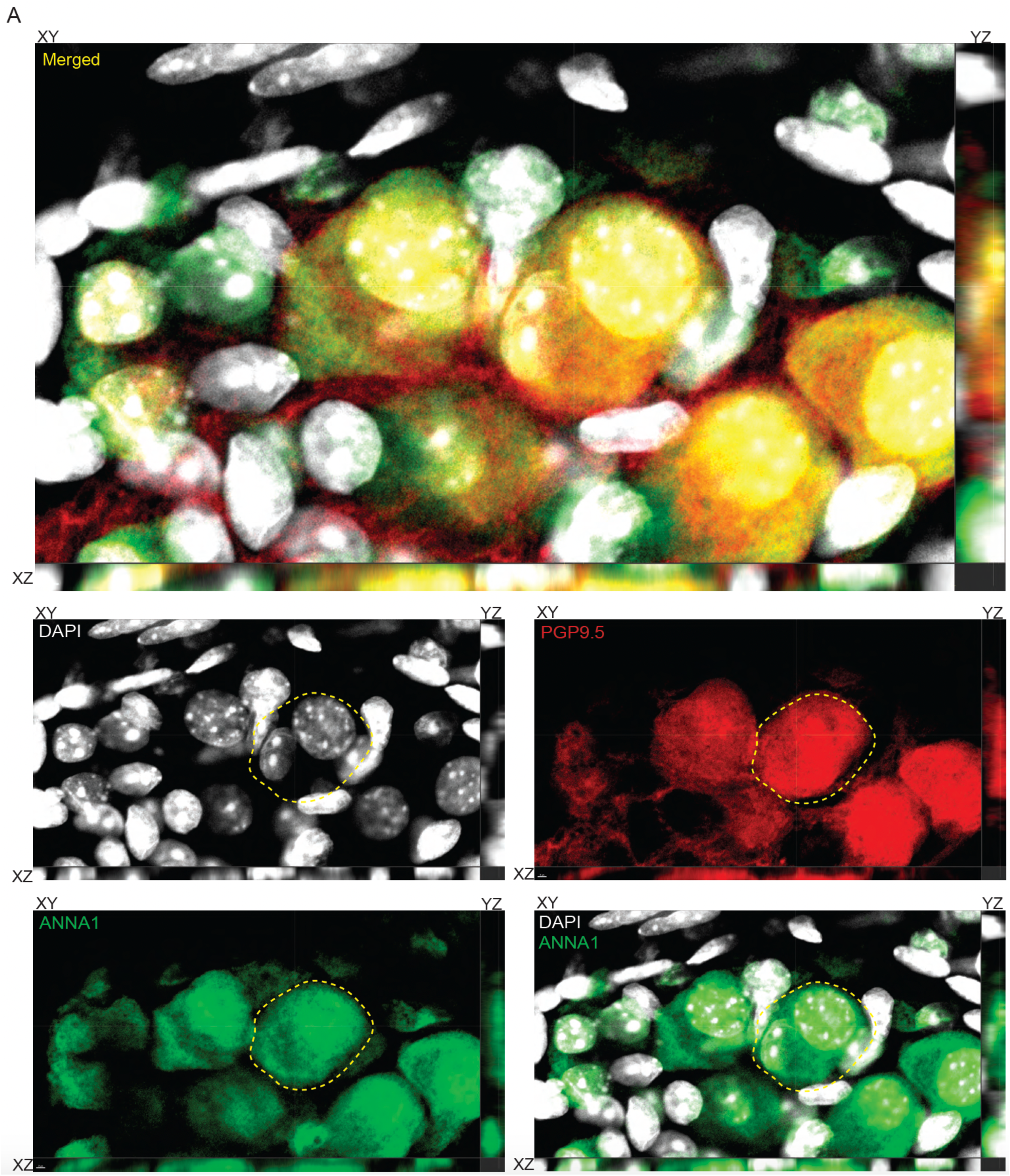
Bi-nucleated ganglionic cells express Hu and PGP9.5. Orthogonal (merged and color-segregated) views of a 3-dimensional confocal microscopy image showing that a DAPI stained (grey), Hu (green) and PGP9.5 (red) immunolabeled cell contains two nuclei. XY, YZ, XY planes of the orthogonal views are denoted for every merged and panel segregated image. Scale bar denotes 2 μm.

### About 10% of all adult murine small intestinal myenteric Hu-expressing cells show evidence of more than 2n DNA content by flow cytometry

While binucleated cells suggest impending cytokinesis in a cell (20), to further study whether cell cycling occurs in Hu-expressing cells, we first used flow cytometry-based estimation of DNA content in adult murine small intestinal Hu-immunolabeled nuclei to test whether myenteric Hu^+^ nuclei show presence of higher than the expected 2N DNA content, which would suggest the presence of cells in S-phase (DNA replication) and in G2/M phase (mitosis) of the cell cycle. On performing this assay on >10,000 unfixed Hu-immunolabeled nuclei derived from LM-MP tissues from adult healthy mice that were housed in the barrier facility at BIDMC, we observed that ∼10% of these Hu^+^ nuclei show evidence of DNA content greater than expected euploidy (Fig 3A, B, C). Based on DNA content, 2.71% of all Hu^+^ nuclei were found to be in S-phase and 7.37% were found to be in G2/M phase of the cell cycle. We next tested whether the observation of >2N DNA content in myenteric Hu^+^ nuclei was dependent on fixation or housing conditions. For this, Hu-immunolabeled fixed nuclei that were derived from small intestinal LM-MP tissue of adult healthy mice housed in a non-barrier facility at Johns Hopkins University were analyzed and were again found to have 2.61% of all Hu^+^ nuclei were found to be in S-phase and 7.39% were found to be in G2/M phase of the cell cycle (Suppl. Fig 2, Fig 3D). Thus, we establish a high degree of concordance in the proportions of Hu^+^ cells in S and G2/M phase of cell cycle in adult mice, irrespective of their housing conditions and colony location, as well as whether the nuclei were stained with or without fixation.

**Figure 3.**
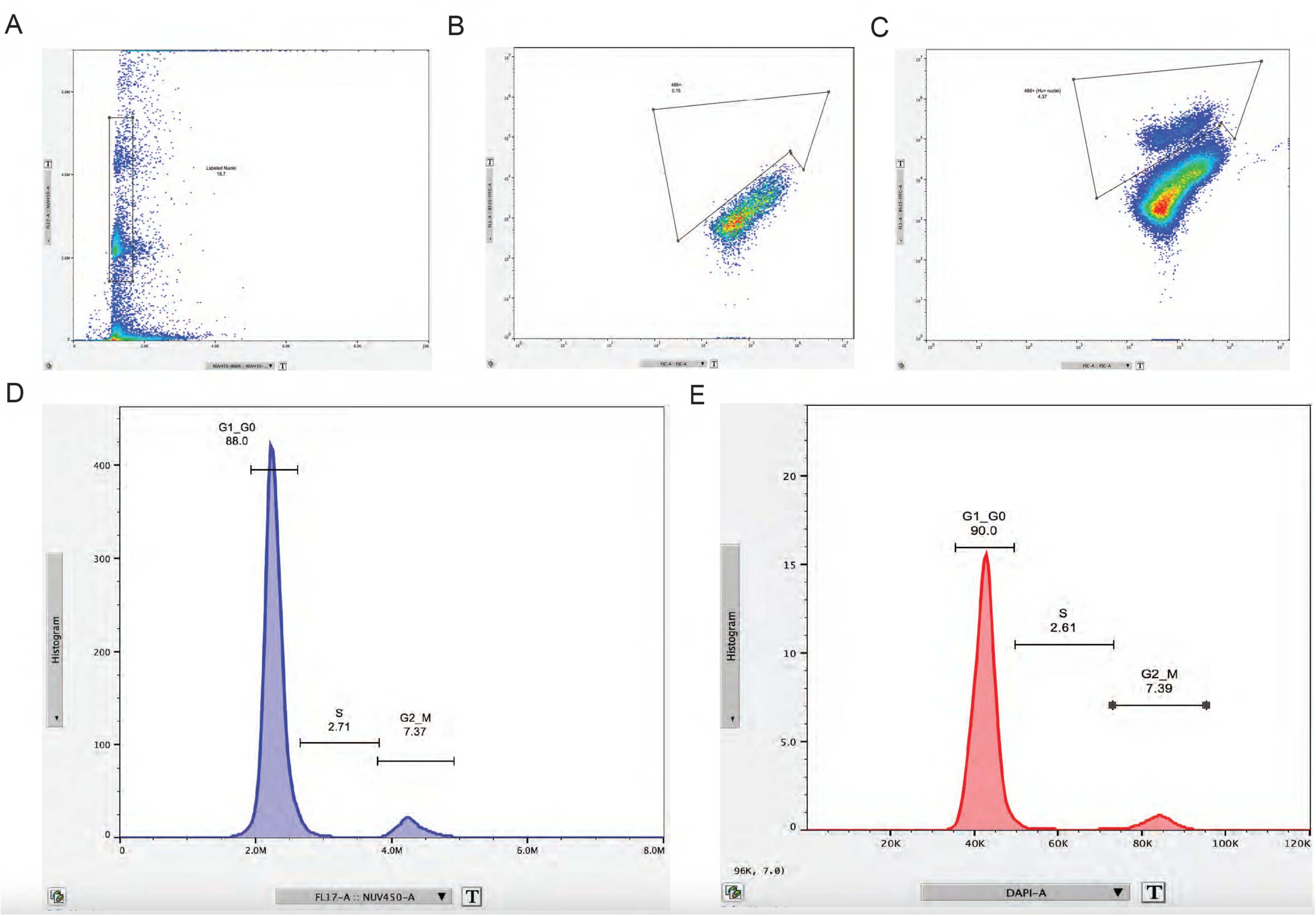
Evidence of S and G2/M phases of cell cycle in a population of Hu-immunolabeled cells from the small intestinal myenteric plexus tissue of adult healthy mice. Flow analyses of nuclei isolated from the Longitudinal muscle and myenteric plexus tissue (LM-MP) from adult healthy mice housed in a barrier facility at BIDMC, when immunolabeled with directly conjugated anti-Hu 488 antibody and co-stained with a DNA-binding dye NucBlue, show that (A) significant proportion of events present with detectable NucBlue-labeling and hence are annotated as nuclei. (B) Using a Hu-unstained population of NucBlue-labeled nuclei, we create gates for Hu-immunolabeled cells, which (C) contain Hu-labeled nuclei in a sample that was immunolabeled with a directly conjugated Hu antibody. (D) This population of isolated Hu- and NucBlue-labeled nuclei from adult murine small intestinal LM-MP tissues from mice from the BIDMC mouse colony, when analyzed using fluorescence intensity of NucBlue as a marker for DNA content, showed that while 88.0% of nuclei have DNA content (2N) expected in the G1 phase cells, 2.71% of Hu^+^ nuclei contain DNA content that corresponds to cells in S-phase (DNA replication phase); and 7.37% of Hu^+^ nuclei contain DNA content that corresponds to cells in G2/M phase (Mitotic phase). The enumeration of nuclei does not include those that show less than 2N DNA content. (E) Flow analysis of nuclei isolated from LM-MP of similarly aged mice housed in at Johns Hopkins University, which were immunolabeled with anti-Hu 647 antibody post-fixation and were stained with the DNA dye DAPI show the presence of 90.0% of nuclei have DNA content (2N) expected in the G1 phase cells, 2.61% of Hu^+^ nuclei contain DNA content that corresponds to cells in S-phase (DNA replication phase); and 7.39% of Hu^+^ nuclei contain DNA content that corresponds to cells in G2/M phase (Mitotic phase).

### About 10% of all adult murine small intestinal myenteric Hu-expressing cells express the cell cycle marker phosphor-Histone H3 (pH3)

In addition to flow analyses of Hu-immunolabeled nuclei, we interrogated whether cycling Hu^+^ cells can be detected in tissues using a known cell cycling marker. For this, we further tested the presence of mitosis in adult enteric Hu^+^ cells by microscopic detection of an established mitotic marker phosphor-histone H3 in the neurons of the myenteric plexus (21–25). By using antibodies against the protein phosphor-Histone H3 (pH3; phosphorylated at residue Ser10, which occurs at the initiation of the G2 phase and continues into metaphase), we observed that the adult healthy small intestinal LM-MP tissue derived from mice housed in the Johns Hopkins University mouse colony contains diverse cell populations of pH3-immunostained cells that are present both within and outside of the myenteric ganglia (Fig 4A, B). Importantly, using the unconjugated pH3 antibody and the anti-Hu ANNA-1 antisera, we observe that the pH3-positive, intra-ganglionic cell population comprises of Hu-immunostained cells (Fig 4A, B), as well as other unlabeled cells that we assume to be populations of proliferating enteric glial cells (26) and enteric neuronal precursor cells, that we have previously shown to express Nestin and Ki-67 (7). Using conjugated pH3 and Hu antibodies, we tested whether adult myenteric ganglia in mice housed in the barrier facility at BIDMC also similarly contained pH3-immunoreactive Hu^+^ cells (Fig 4C). Similar to our earlier observations (7), Cleaved Caspase 3 (CC3) immunoreactivity, which indicates apoptotic cells, was found in a subset of myenteric Hu^+^ cells (Fig 4D). We further tested whether pH3 immunoreactivity was found both in adult murine small intestinal enteric neural precursor cells (ENPC) that express Nestin but do not express Hu, and in ganglionic cells that co-express Nestin and Hu. By co-immunolabeling, we find that pH3 immunolabeling was found both in Nestin-expressing ENPC, as well as in Hu-expressing cells that show detectable but low expression of Nestin (Fig 5, 6). This suggests that Hu^+^ pH3^+^ ganglionic cells are distinct from Nestin-expressing cycling ENPC.

**Figure 4:**
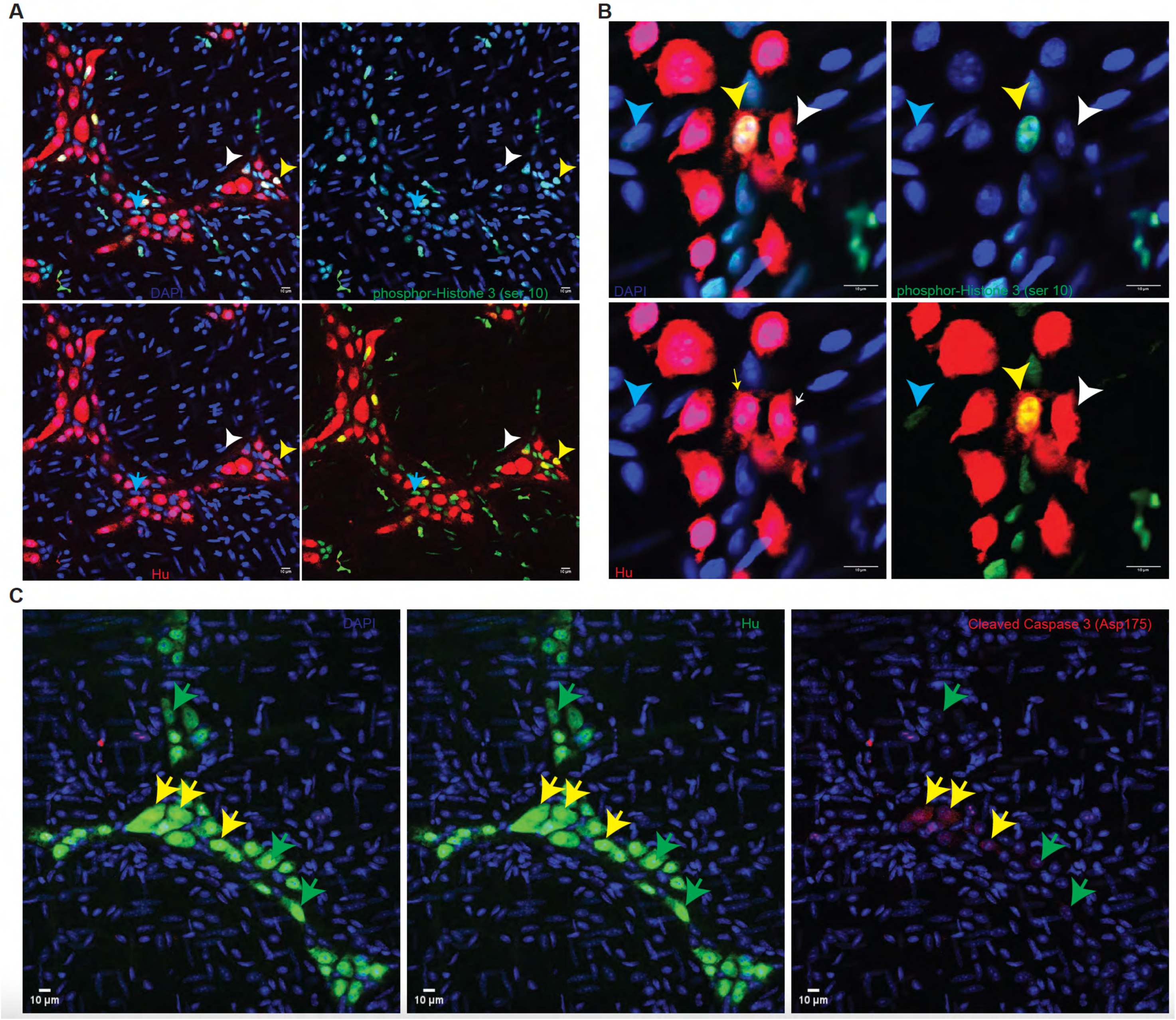
Cell proliferation marker phosphor-histone H3 (pH3) is detected in neurons and other cells of the adult healthy myenteric plexus tissue. Confocal microscopy shows that the (A) Longitudinal muscle and myenteric plexus tissue (LM-MP) from an adult healthy mouse that was housed at Johns Hopkins University, when stained with unconjugated antibodies against phospho-Histone H3 (pH3, red) and the pan-neuronal marker Hu (green) shows the presence of pH3 immunoreactivity in ganglionic cells that do express Hu (yellow arrow) and those that don’t express Hu (cyan arrow). pH3 immunoreactivity is also detected in extra-ganglionic cells in this tissue (white arrow) that are presumed to be longitudinal muscle cells. (B) Magnified image of a myenteric ganglia labeled with antibodies against pH3 (red) and Hu (green) again show the presence of pH3-immunoreactive Hu-immunolabeled new-born neurons (yellow arrow), along with neurons that are not immunoreactive against pH3 (white arrow) and pH3-immunoreactive extra-ganglionic cells (cyan arrow). (C) *Left:* Representative image of LM-MP tissue from an adult mouse housed in the barrier facility at BIDMC when stained with directly conjugated antibody against pH3 (red) shows the presence of significant pH3-immunoreactivity in the tissue. The tissue was also co-immunostained with directly conjugated anti-Hu antibody and the region of interest (white box) when magnified shows (*Right*) the presence of Hu-immunolabeled (green) neurons that co-label for pH3. Nuclei are stained with DAPI (blue). (D) Representative image of the adult murine small intestinal LM-MP tissue, when stained with antibodies against Cleaved Caspase 3 (CC3, red) and the pan-neuronal marker Hu (green), shows the presence of CC3 immunoreactivity in a subset of Hu-immunoreactive neurons (yellow arrows), while other neurons in the same and other ganglia do not immunolabel for CC3 (green arrow). Nuclei are labeled with DAPI (blue). Scale bar indicates 10 µm.

**Figure 5:**
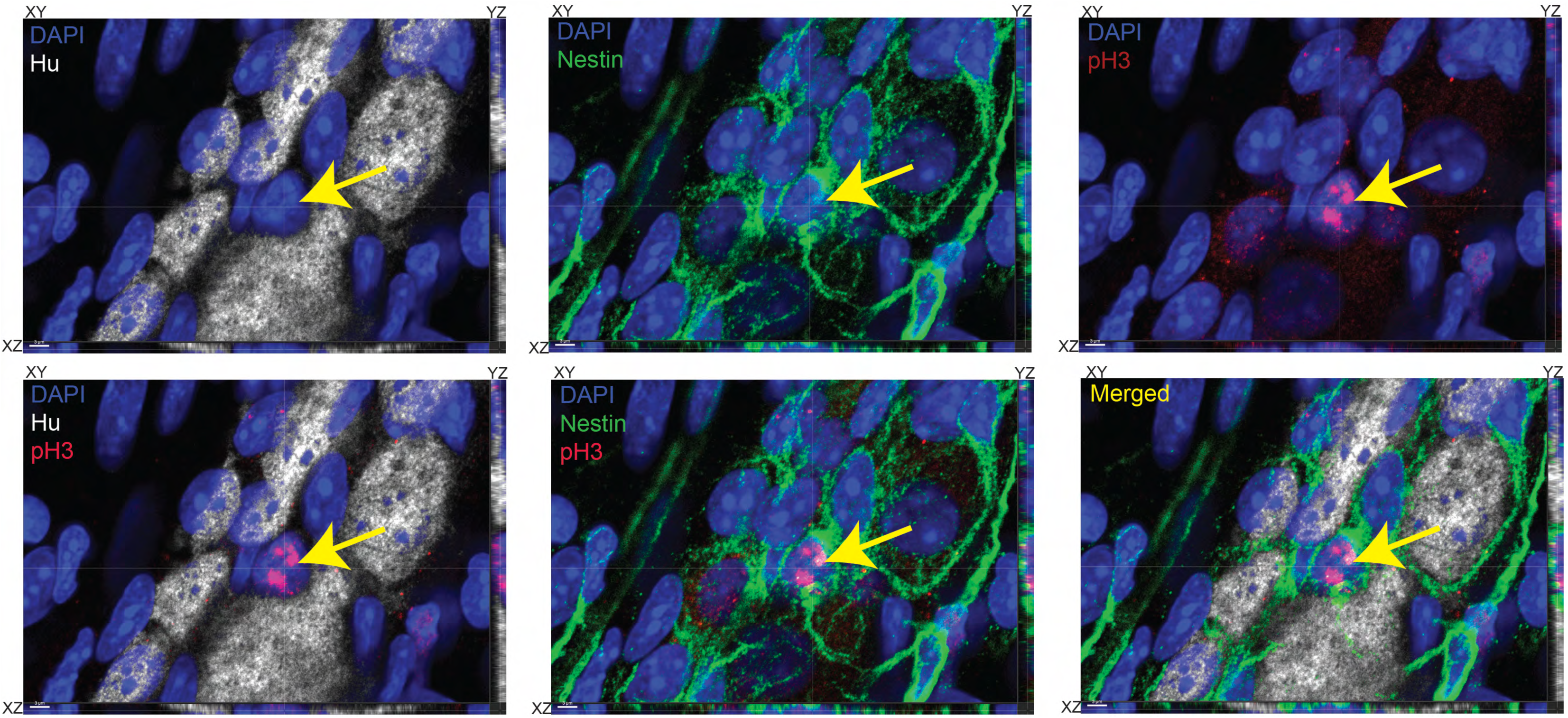
Nestin-expressing and Hu-non-expressing enteric neural precursor cells express cell cycle marker phosphor-Histone H3. Orthogonal (merged and color-segregated) views of a 3-dimensional confocal microscopy image showing a myenteric ganglion from an adult murine small intestinal tissue, where the tissue is immunolabeled with antibodies against Hu (grey), phosphor-Histone H3 (pH3, red), Nestin (green), and is stained with nuclear dye DAPI (blue). Yellow arrow points out a Nestin and pH3-immunolabeled cell that does not immunolabel for Hu. Presence of pH3 immunoreactivity and absence of Hu immunoreactivity in this Nestin-immunolabeled cell suggests it is a cycling enteric neural precursor cell. XY, YZ, XY planes of the orthogonal views are denoted for every merged and panel segregated image. Scale bar denotes 3 μm.

**Figure 6:**
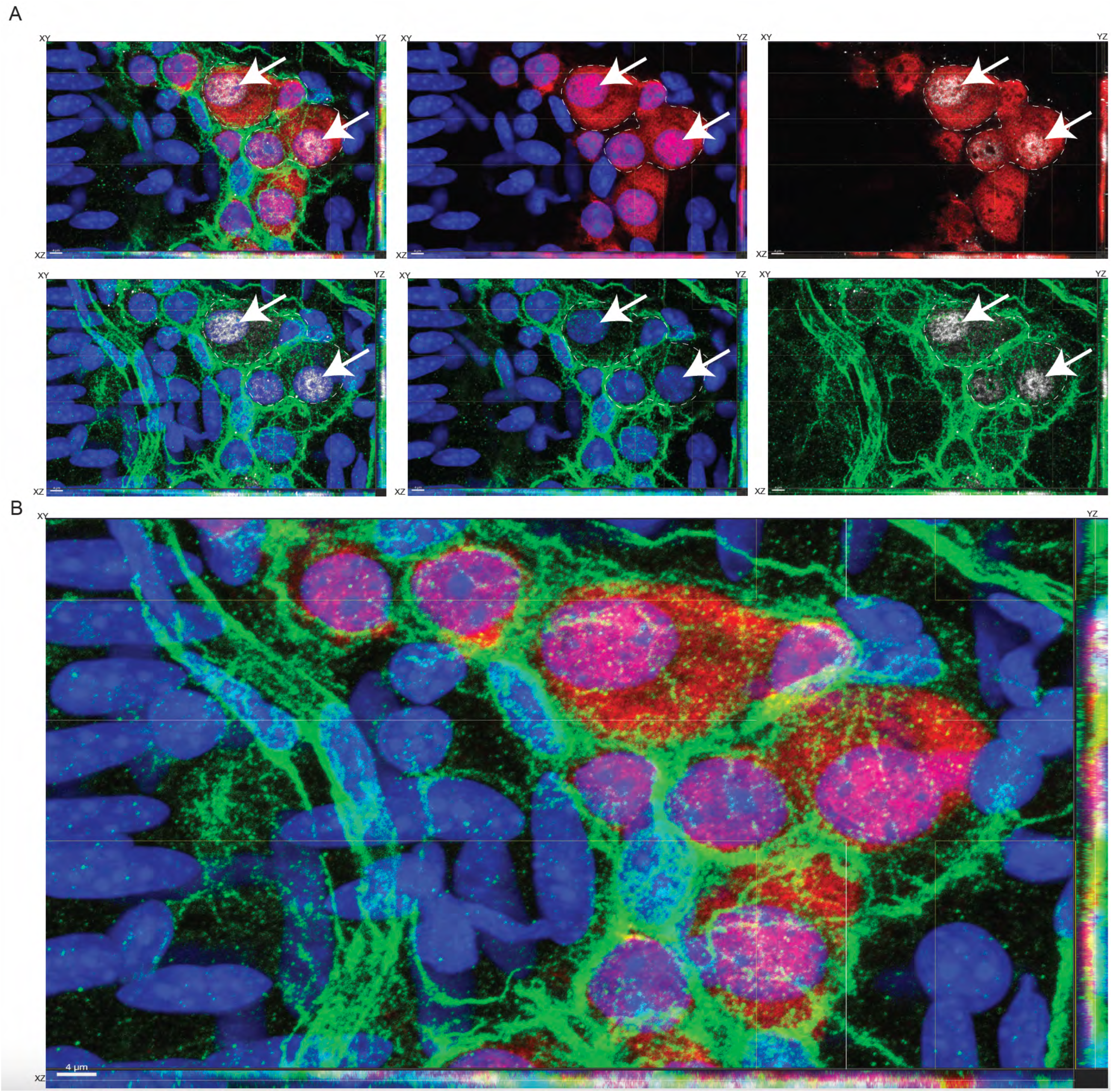
Hu-expressing cells that express cell cycle marker pH3 also exhibit Nestin immunoreactivity. Orthogonal (A) color-segregated and (B) color-merged image of a myenteric ganglion from an adult murine small intestinal tissue, where the tissue is immunolabeled with antibodies against Hu (red), phosphor-Histone H3 (pH3, grey), Nestin (green), and is stained with nuclear dye DAPI (blue). Dashed white lines depict a multi-nucleated contiguous Hu-immunolabeled cell, with white arrows showing that two of the three nuclei are immunolabeled with antibodies against pH3. The pH3^+^ Hu^+^ cells (white arrows) also show positive immunostaining for Nestin. XY, YZ, XY planes of the orthogonal views are denoted for every merged and color-segregated image. Scale bar denotes 4 μm.

We have previously reasoned that isolated adult murine small intestinal LM-MP tissues require gentle fixation (12, 18). To test whether the presence of pH3 in Hu^+^ ganglionic cells is an artifact of gentle fixation, we tested three additional timepoints of 10, 15, and 20 minutes of fixation and found that pH3 immunoreactivity was present in Hu-immunolabeled cells at each of these timepoints. Similar to our observations on binucleated Hu^+^ cells in gently fixed tissues, tissues fixed for 20 minutes again showed the presence of binucleated Hu+ cells that showed evidence of asymmetric cell division (Fig 7).

**Figure 7:**
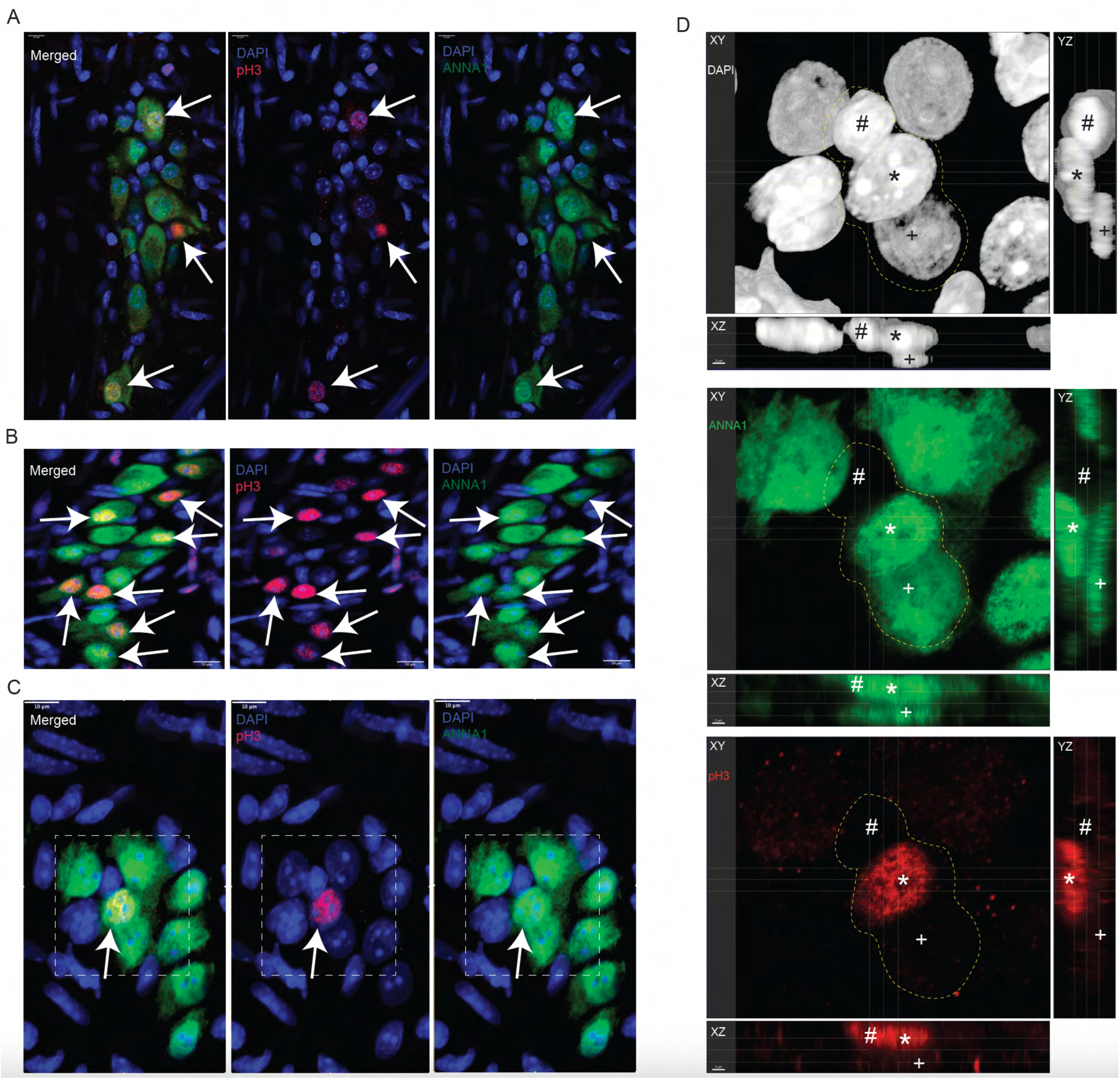
pH3-immunoreactivity in Hu^+^ cells is unaffected by fixation conditions. Immunostaining adult murine small intestinal LM-MP tissues that were fixed with 4% paraformaldehyde for (A) 10 minutes, (B) 15 minutes, and (C) 20 minutes and subsequently immunostained with anti-Hu antibodies in ANNA1 serum (green) and pH3 (red) and counterstained with DAPI (blue). Merged and color segregated views are shown for each of the treatments. White arrows show the presence of pH3-immunoreactivity in the DAPI stained nuclei of Hu-immunoreactive cells in each of the treatments. White square in panel (C) is magnified in (D) and viewed in orthogonal views where XY, YZ, and XZ planes are shown. A contiguous nuclear structure (stained with DAPI, grey) containing 3 nuclear lobes ‘#, *, +’ are observed within dashed white lines. The nuclear lobe ‘#’ is unstained by pH3 (red) and ANNA1 (green) antibodies, lobe ‘*’ is stained by both pH3 and ANNA1, and lobe ‘+’ is stained by ANNA1 but not by pH3. The three panels show color segregated views depicting an asymmetric cellular structure where parts of the nuclear structure exhibit Hu and/or pH3 immunoreactivity, while another part does not. XY, YZ, XY planes of the orthogonal views are denoted for every merged and color-segregated image. Scale bar in (A – C) denotes 10 μm and in (D) denotes 2 μm.

Since co-immunolabeling with antibodies against pH3 and pan-neuronal marker Hu provides evidence of mitosis in adult small intestinal myenteric Hu^+^ cells, we next quantified the proportions of pH3-immunoreactive mitotic small intestinal myenteric Hu^+^ cells at steady state in adult mice. By confocal microscopy, we observed 2577 Hu-immunolabeled Hu^+^ cells from the small intestine of 6 adult healthy male and female mice (n = 3 of either sex), which included myenteric plexus tissue from both duodenum and ileum. We found that 8.54% ± 1.09 (mean ± SEM) of all Hu-immunolabeled cells also showed co-immunolabeling with anti-pH3 antibodies (Fig 8 A,B).

**Figure 8:**
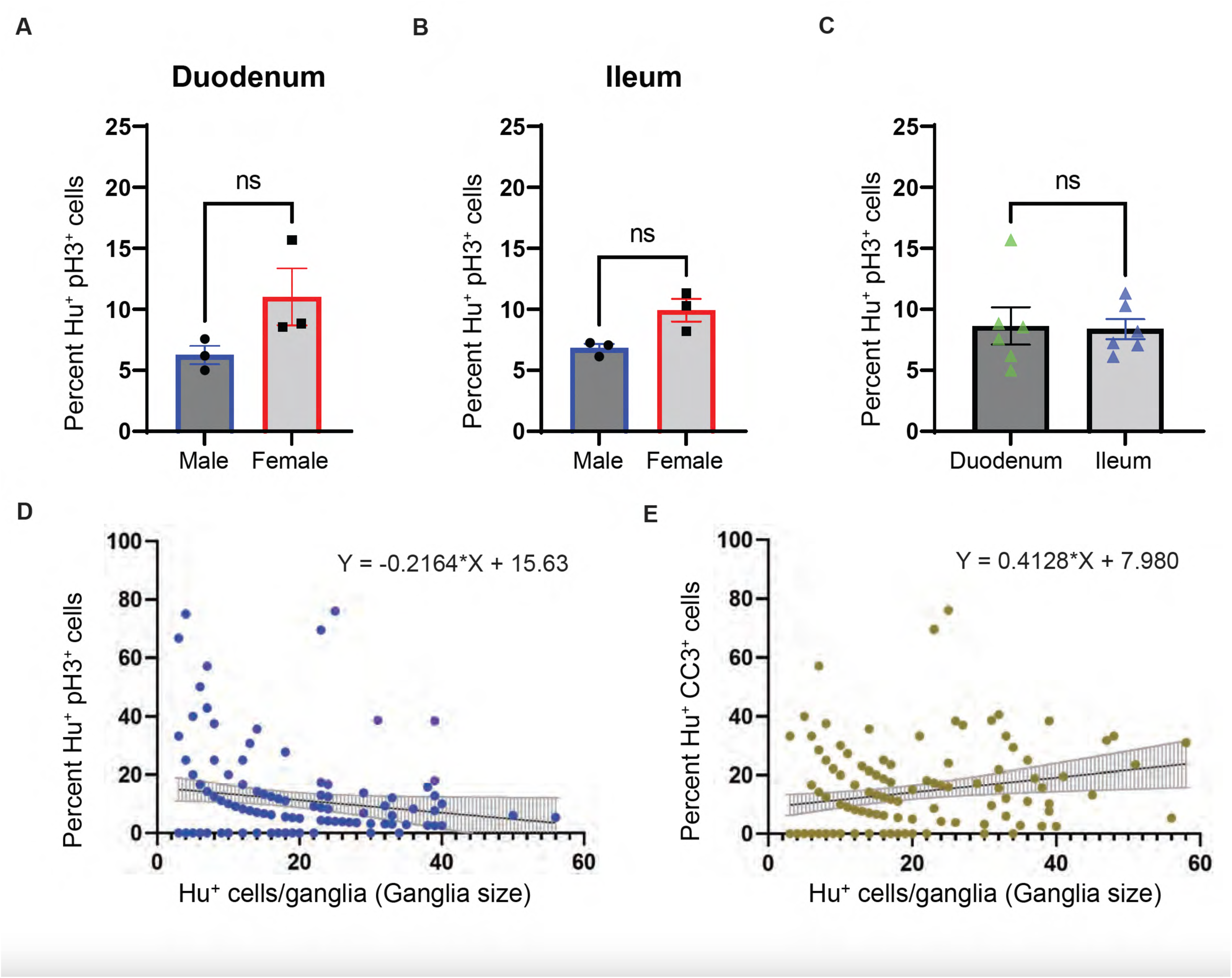
Quantifications of pH3-immunoreactive and CC3-immunoreactive Hu^+^ cells in the adult murine gut. (A) Quantification of pH3-immunoreactive Hu^+^ cells in the duodenal tissue of age-matched male and female mice show no significant sex-bias in their proportions (p > 0.05, Students’ t-test). (B) Quantification of pH3-immunoreactive Hu^+^ cells in the ileal tissue of age-matched male and female mice show no significant sex-bias in their proportions (p > 0.05, Students’ t-test). (C) Quantification of pH3-immunoreactive Hu^+^ cells in the duodenal and ileal tissue of age-matched mice show no significant differences in their proportions between the two tissue regions (p > 0.05, Students’ t-test). (D) Distribution of percent pH3-immunoreactive Hu^+^ cells in small intestinal ganglia containing various numbers of Hu^+^ cells. (E) Cubic curve fit (black line) for the distribution of CC3-immunoreactive Hu^+^ cells in small intestinal ganglia containing various numbers of Hu^+^ cells.

Next, we analyzed whether pH3 immunoreactivity in Hu^+^ cells from ileal or duodenal myenteric ganglia showed any sex bias. In the duodenal myenteric ganglia, we counted 670 Hu^+^ cells in tissues from female mice and 601 Hu^+^ cells in the tissues from male mice (n = 3 of either sex), and found no significant difference between the percentage of pH3-immunoreactive Hu^+^ cells between the two sexes (mean ± SEM pH3^+^ neurons: female mice = 11.63 ± 2.00; male mice = 6.26 ± 0.75; p = 0.17, Unpaired t-test with Welch’s correction, Fig 8 A). In myenteric ganglia of the ileum, we counted 667 Hu^+^ cells in the tissues from female mice and 639 Hu^+^ cells in the tissues from male mice (n = 3 of either sex), and found no significant difference between the percentage of pH3-immunoreactive Hu^+^ cells between the two sexes (mean ± SEM pH3^+^ neurons: female mice = 9.93 ± 0.92; male mice = 6.82 ± 0.34; p = 0.06, Unpaired t-test with Welch’s correction, Fig 8 B). Finally, we compared pH3 immunoreactivity in populations of Hu^+^ cells between ileal and duodenal tissue across the two sexes and found no significant difference (Duodenal tissue: Hu^+^ cells counted = 1271, mean ± SEM pH3^+^ Hu^+^ cells: 8.64 ± 1.52; Ileal tissue: Hu^+^ cells counted = 1306, mean ± SEM pH3^+^ Hu^+^ cells: 8.38 ± 0.82; p = 0.88, Unpaired t-test with Welch’s correction, Fig 8 C).

We have previously shown that there is significant heterogeneity in ganglia size, demonstrated by small intestinal myenteric ganglia containing as few as 3 neurons or over 100 neurons (13). Here, we tested whether the proportion of mitotic Hu^+^ cells correlate with ganglia size in the adult healthy small intestinal myenteric plexus. By analyzing the proportions of pH3-immunoreactive Hu^+^ cells in 147 ganglia of various sizes (3 – 56 neurons per ganglia) from duodenum and ileum from 6 mice together, we found an absence of any significant correlation between percent Hu^+^ cells immunoreactive for pH3 in myenteric ganglia and the numbers of Hu^+^ cells within ganglia (ganglia size) (D’Agonistino and Pearson test for normality showed non-normal distribution, non-parametric Spearman correlation test: r = −0.05625 p = 0.5; Fig 8 D).

We next tested whether there was any correlation between ganglia size and the proportions of CC3-immunoreactive apoptotic Hu^+^ cells in the adult healthy murine small intestinal myenteric plexus. We quantified the percent of CC3-immunoreactive Hu^+^ cells from 83 myenteric ganglia from 3 mice, where the myenteric ganglia size ranged from 3 – 58 neurons. We found that the proportion of CC3^+^ Hu^+^ cells in small intestinal myenteric ganglia increased significantly with ganglia size (D’Agonistino and Pearson test for normality showed non-normal distribution, non-parametric Spearman correlation test: r = 0.4181 p <0.0001; Fig 8 E).

### Human small intestinal myenteric ganglia show evidence of neuronal turnover

We next tested whether the adult human small intestinal myenteric plexus tissue also shows evidence of significant cell proliferation through the presence of pH3-immunoreactive Hu^+^ cells. In immunolabeled FFPE tissue sections, by using directly conjugated antibody against pH3, we found that many cells in the longitudinal muscle layer of human gut wall also showed immunoreactivity against the mitosis marker pH3 (Fig 9 A). By co-immunostaining these FFPE tissue sections with antibodies against pH3 and Hu, we found that many pH3 immunoreactive cells were found in the myenteric ganglia, some of which were also immmunolabeled by antibodies against Hu (Fig 9 B). We enumerated proportions of pH3^+^ Hu^+^ cells in myenteric ganglia in FFPE tissue sections from duodenal full thickness tissues derived from 4 human samples. We counted a total of 152 Hu^+^ ganglionic cells (mean ± standard error of Hu^+^ cells counted per sample = 37.5 ± 9.56) and found that 23.43 ± 5.22 (mean ± S.E) percent of Hu^+^ cells also immunolabeled for the mitosis marker pH3. This data provides evidence that a significant proportions of human Hu^+^ myenteric ganglia cells are mitotic in nature.

**Figure 9:**
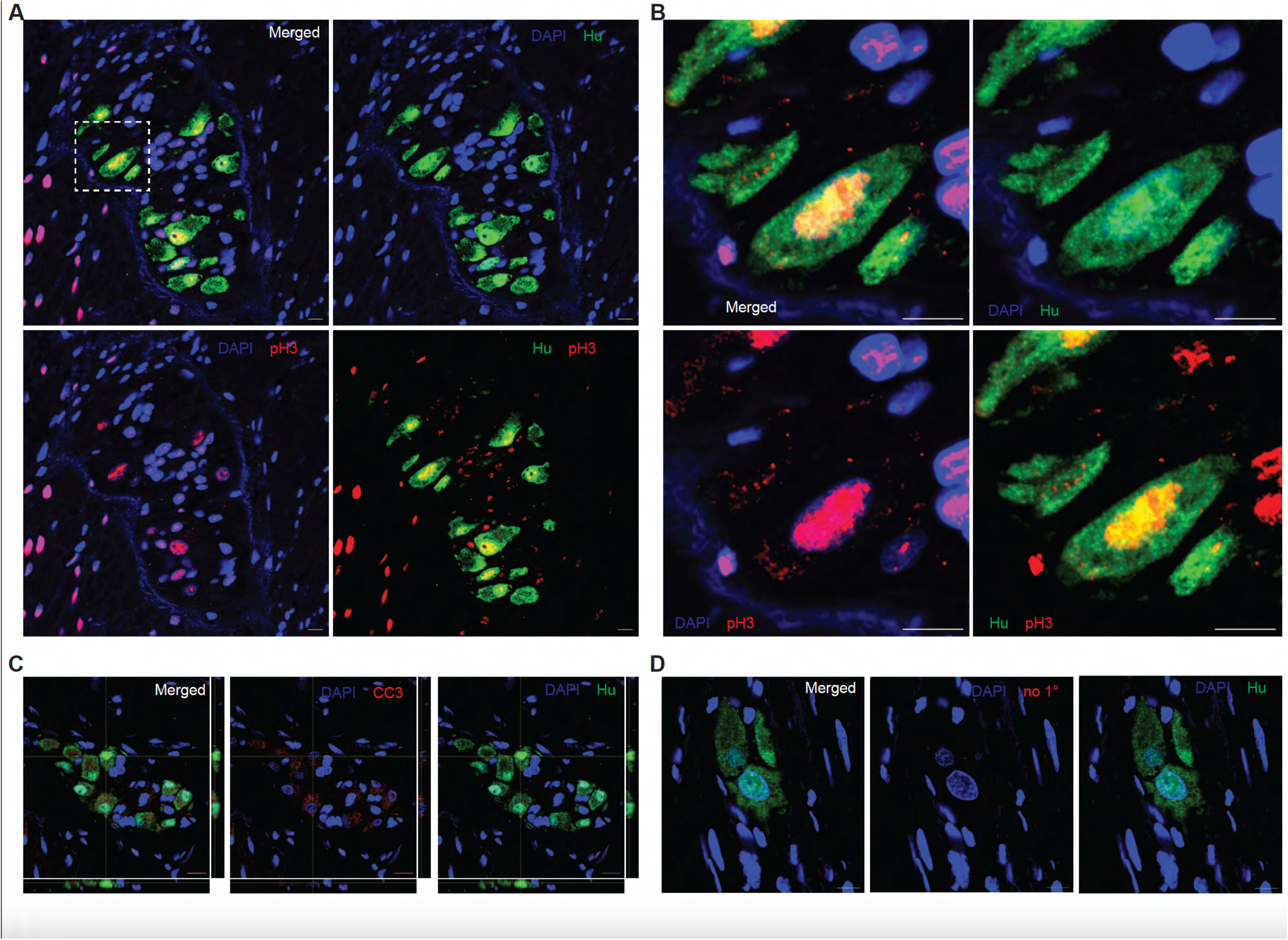
Hu^+^ myenteric cells expressing cell cycle marker pH3 or expressing apoptotic marker Cleaved Caspase 3 exist in human small intestinal myenteric ganglia. (A) Cells of the longitudinal muscle layer of the human gut wall in formalin fixed paraffin embedded (FFPE) tissue sections from adult human small intestine with immunolabelling of mitotic marker pH3 (red) and co-staining with DAPI (blue) show detectable pH3 immunoreactivity in many cells of this tissue. Image created by stitching contiguous 40X images of the section which was imaged with the EVOS M7000 microscope. (B) Co-immunolabeling adjacent sections of this tissue with directly conjugated antibodies against mitotic marker pH3 (red) and pan-neuronal marker Hu (green) and by imaging them with the M7000 microscope, we observe the presence of pH3-immunoreactive cells that also immunolabel with antibodies against Hu (yellow arrows). (C) Confocal microscopy of FFPE human tissues that were immunolabeled with anti-Hu ANNA-1 antisera and anti-pH3 antibody and then suitably co-stained with secondary antibodies again show the presence of pH3 and Hu co-labeled cells within the myenteric ganglia (yellow arrow). (D) Orthogonal views generated from confocal microscopy of a myenteric ganglia immunostained with antibodies against CC3 (red) and anti-Hu ANNA-1 (green) show the presence of CC3-immunolabeled neurons. (E) Representative image of a myenteric ganglia immunostained with antibodies against Hu (green) but no primary antibodies for the red channel shows a lack of non-specific staining with the secondary antibodies. Photomicrographs in panels (C – E) were generated by confocal microscopy. Nuclei are labeled with DAPI (blue) and scale bar in (C – E) indicates 10 µm.

Similarly, we also found that the apoptotic marker Cleaved Caspase 3 was detected in a subset of human small intestinal myenteric neurons (Fig 9C), while the negative control for CC3 immunoreactivity (performed without primary antibody) showed no background signal (Fig 9D).

## Discussion

Using established microscopic, flow cytometric, and immunolabeling based assays (21–25, 27–29), our report provides evidence that a significant proportion of Hu^+^ ganglionic cells in the adult small intestinal myenteric plexus tissue undergo mitosis. Neurogenesis from precursor cells requires DNA replication and mitosis, and the detection of mitosis and apoptosis in Hu-immunolabeled cells in healthy small intestinal myenteric plexus tissue shows that there is ongoing genesis of neurons in this organ, which offsets the continual loss of neurons to apoptosis. Our prior study, which used thymidine analogue-based assays to demonstrate that almost 90% of adult small intestinal neurons were generated during a 2-week period, had suggested that ∼10% of neurons are generated (and hence are newborn) on any given day(7). Here, our DNA content-based flow cytometry analysis shows that ∼10% of Hu-immunolabeled myenteric nuclei showed evidence of DNA content associated with G2/M (or mitotic) phase of the cell cycle. This is consistent with our microscopy-based data that demonstrate the presence of binucleate Hu^+^ cells and the immunolabeling-based data that show that ∼10% of Hu-immunolabeled cells within myenteric ganglia express the mitotic marker phosphor-histone H3 (pH3). Thus, our observation of ongoing mitosis in Hu-immunolabeled cells using these different methods closely matches the rate of neurogenesis we had previously deduced using detection of thymidine analogues (7). In addition, the presence of pH3 can be readily detected in >20% of Hu-immunolabeled ganglionic cells in the adult human gut – further suggesting that a similar neurogenic process is also present in the adult human gut. We again confirm that a subpopulation of adult murine small intestinal myenteric neurons showed detectable presence of the apoptotic marker Cleaved Caspase 3 (CC3) (7).

Evidence of DNA replication (S-phase) and mitosis (G2/M phase) in Hu-immunolabeled cells suggests that in addition to being a marker for mature functional neurons, Hu is also expressed by cycling neuroblasts or transit amplifying cells that are committed to a neuronal fate. Thus, rather than suggesting that adult enteric neurons are mitotic in nature, we infer that Hu immunoreactivity is not restricted to mature neurons. This argument is supported by prior studies that have shown that Hu is expressed in neuronally committed cycling progenitors or neuroblasts as well as in immature neurons (30, 31). The presence of a significant population of cycling Hu^+^ neuroblasts in the adult myenteric plexus provides evidence of a robust steady state neurogenic process in the adult gut.

The thymidine analogue-based assay we had previously deployed to observe adult enteric neurogenesis requires complicated methods (12). In addition, by virtue of being based on the uptake, incorporation, and accumulation of these chemicals over time into newly synthesized nucleic acids, this assay can neither sensitively measure the ongoing rate of DNA synthesis and cell proliferation at any given point of time, nor can it be used to assess rate of neurogenesis in humans. With flow analyses, immunolabeling and microscopy-based methods, we use simple established assays to robustly measure the proportions of cycling neuroblasts in tissue to infer the rate of neurogenesis at health in the adult human and murine gut.

Furthermore, two previous studies suggested that the proportions of apoptotic myenteric neurons are significantly greater in the human gut than in the murine gut (human gut 20 – 30% apoptotic neurons vs 10 – 11% in murine gut) (4, 5, 19). Thus, it would be expected that the rate of genesis of neurons in the adult human gut would also be greater than in murine gut. Our results, which show that ∼23 - 24% of all human myenteric Hu^+^ cells immunolabel for the mitotic marker pH3, closely match the published rate of apoptosis in human ENS neurons. These data demonstrate that as in the murine gut, the proportions of newborn and apoptotic neurons in the human gut are evenly matched. This suggests that as established by us previously in mice, continual neurogenesis and neuronal loss are also a feature of the adult human ENS.

The immunofluorescence analyses of mitotic neuroblasts and apoptotic neurons in the murine small intestinal myenteric plexus also allowed us to test whether neurogenesis and neuronal loss - the two processes driving neuronal turnover - occur equally in all myenteric ganglia across two different small intestinal regions and between the two sexes. Taking the proportions of mitotic neuroblasts to mature neurons as a measure of the neurogenic activity for each ganglion, we provide evidence that neurogenic activity does not significantly differ between differently sized ganglia, between two different intestinal regions, or between sexes. However, while the rate of neurogenesis does not change across ganglia, we observe that larger ganglia (i.e. with greater number of neurons) have a higher percentage of neurons undergoing apoptosis. If neurogenesis occurs at a uniform rate across all ganglia, then the observation that apoptosis occurs more frequently in larger ganglia implies that there may be a regulatory process that discourages the preservation or further enlargement of these larger ganglia. This is reflected in our prior findings that showed that smaller ganglia are much more frequent than larger ganglia in the adult small intestinal myenteric plexus tissue (13). This suggests that the balance between the genesis and death of neurons within ganglia is crucial for regulating ganglia size. Furthermore, it would indicate that any disruptions to this balance, whether through increased neurogenesis or elevated neuronal death, could alter the normal proportions of smaller and larger ganglia, potentially leading to disorders in intestinal movement. While whether such a ganglia size-associated skew in genesis and loss of neurons extends to human tissue is still unknown, it was recently demonstrated (32) that patients with dysmotility have significantly reduced neuronal numbers in myenteric ganglia (or that the presence of larger ganglia in their ENS is greatly reduced). These are similar to other observations (33) that observed fewer numbers of myenteric neurons in patients with ulcerative colitis, than in healthy controls. These observations suggest that ganglionic rate of neurogenesis and of neuronal loss may change in disease to alter the ENS structure.

That adult enteric neurogenesis at steady state occurs in the adult ENS has been controversial (34). Despite our prior study that observed significant proportions of label-retaining myenteric neurons in mice dosed with thymidine analogues (7), other investigators were unable to replicate these findings and detect thymidine analogue-based label-retention in neurons (35). While it may be argued that the inability to detect enteric neurogenesis at steady state in the adult healthy gut may be due to technical challenges and issues (36), our study furthers the case for steady state adult enteric neurogenesis by testing the presence of DNA synthesis and mitosis in Hu-expressing myenteric cells. A recent report studied adult enteric neurogenesis at health and in disease using a different immunohistochemical marker for new-born neurons Sox2, which was also found to label ∼10% of all Hu^+^ myenteric cells in the adult healthy colon (37). The concordance between the proportions of Sox2^+^ Hu^+^ ‘newborn’ neurons in the colon in their study, and the proportions of pH3^+^ Hu^+^ myenteric cells and Hu^+^ cells in G2/M phase in our study provides evidence that myenteric neurons in the small intestine and colon do turn over in significant numbers, that the rate of neurogenesis or proportions of mitotic neuroblasts matches the observed rate of neuronal loss across various studies (6, 8, 9, 38), and that steady state neurogenesis is an important mechanism through which adult ENS homeostasis is maintained (12).

While this study furthers our model of continual neurogenesis to support neuronal homeostasis in the adult ENS, further work is needed to answer important questions that emerge considering this data and our other recent reports. Our recent study provides evidence that the adult ENS consists of equal proportions of neurons derived from neural crest and mesodermal lineages (38), and our prior study using thymidine analogues found that ∼90% of adult enteric neurons turnover during a 2-week period (7). The near comprehensive nature of neuronal turnover in the adult ENS suggests that in addition to genesis of neural crest (NC)-derived enteric neurons (NENs), which we showed occurs from Nestin-expressing NC-derived precursor cells, the population of mesoderm-derived enteric neurons (MENs) also turn over at steady state. While our prior report found molecular evidence of continual genesis of MENs in the post-natal ENS (18), this current report does not test the identity of precursor cells for the MEN lineage. We propose that populations of both neural crest-derived enteric neurons (NENs) and the mesoderm-derived enteric neurons (MENs) undergo steady state neurogenesis in adults. This suggests that there are two separate lineage-associated populations of enteric neuronal precursor cells in the adult ENS – one for NENs (that co-express Nestin and p75NTR), and another for MENs, whose exact molecular identity currently remains unknown (38). The data presented in this report, taken in the context of our prior reports, suggests that pH3^+^ Hu^+^ cells or Hu^+^ cells that are in G2/M phase include neuroblasts for both MENs and NENs. Future work will focus on establishing how ganglionic Hu^+^ cells of the mesodermal and NC-lineage differ in their proportions of cycling cells at different ages. In addition, our microscopic assessment of myenteric ganglia provide evidence of binucleated cells, and of pH3-immunolabeled Hu^+^ cells that appear to undergo asymmetric division into Hu^−/+^ cells. Mechanisms of cell division include canonical nucleokinesis and cytokinesis, as well as non-canonical endoreduplication and closed mitosis, through which cells may increase their nuclear DNA content without cytokinesis or maintain their nuclear membranes while undergoing cytokinesis (22, 39, 40). While the presence of asymmetric Hu-immunolabeled cells, where cell structure shows varying immunolabeling for Hu and pH3, suggests the presence of non-canonical cytokinetic mechanisms in the adult ENS, the nature of the precise cell biological mechanisms through which Hu^+^ cells cycle need to be further interrogated in detail. This study provides the rationale for studying these mechanisms in future.

Our study is robust as it establishes concordance of our observations through different approaches - the high resolution microscopy approach to test the presence of bi-nucleated Hu-expressing cells, DNA content-based flow analyses to test the presence of diploid DNA content indicative of S- and G2/M phase, and the pH3-based immunofluorescence-assay based approach to identify and enumerate the proportions of mitotic Hu-expressing cells in adult murine and human ENS. Thus, this study further confirms the presence of the homeostatic mechanism of steady state neuronal turnover in the adult ENS.

## Acknowledgements

This work was supported by funding from NIA R01AG66768, R21AG072107, Diacomp Foundation (Pilot award Augusta University) and Pilot grant from the Harvard Digestive Disease Core (SK). AG was supported by a Fulbright Future Scholarship, funded by The Kinghorn Foundation. J.S. was funded through the Maryland Genetics, Epidemiology, and Medicine training program sponsored by the Burroughs Welcome Fund and from Walter Benjamin Fellowship (528835020) from Deutsche Forschungsgemeinschaft (PS). We acknowledge the help of Mr. John Tigges, Technical Director/Manager Flow Cytometry Science Center, Beth Israel Deaconess Medical Center for his help with flow cytometric analyses on Flowjo. We appreciate the help of Dr. Taru Muranen and Dr. Nina Kozlova for their help with Flowjo.

---

**Supplementary Figure 1: Observation of bi-nucleated or conjoined nuclei in Hu^+^ myenteric cells.** Orthogonal color-merged and color-segregated views of an image of a myenteric ganglion from an adult murine small intestinal tissue, where the tissue is immunolabeled with antibodies against Hu (green) and stained with nuclear dye DAPI (grey) shows the presence of two near or conjoined nuclei # and + in a contiguous Hu-immunolabeled cell. Scale bar denotes 1 µm.

**Supplementary Figure 2: Flow gates used for assessing Hu-immunolabeled nuclei from adult small intestinal LM-MP tissues from mice in the Johns Hopkins colony.** Fixed nuclei isolated from adult murine small intestinal LM-MP were stained with nuclear dye DAPI and directly conjugated Hu antibody (Alexa 647) and assessed to establish the gates for DAPI+ nuclei (left plot), and DAPI+ nuclei that immunolabeled for Hu (right plot).

