## Supplementary figures and images for "Detection of mitotic neuroblasts provides additional evidence of steady state neurogenesis in the adult small intestinal myenteric plexus"

### Suppl. Fig 1

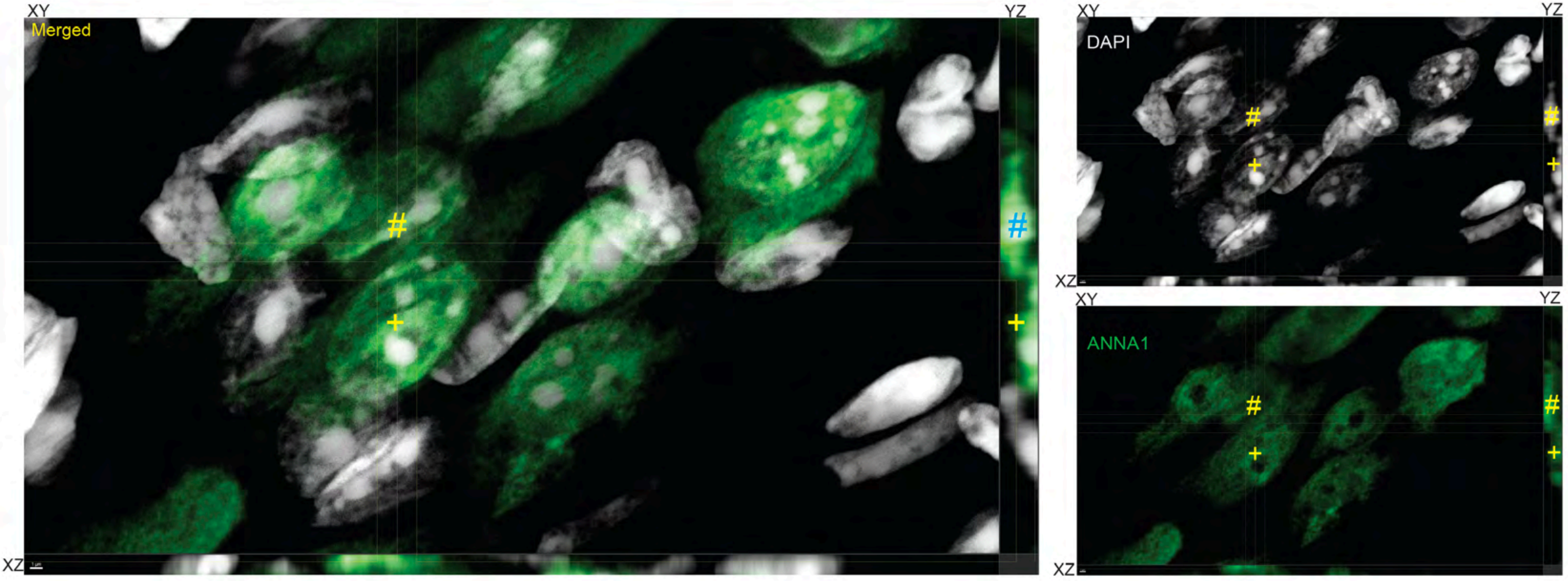

### Suppl. Fig 2

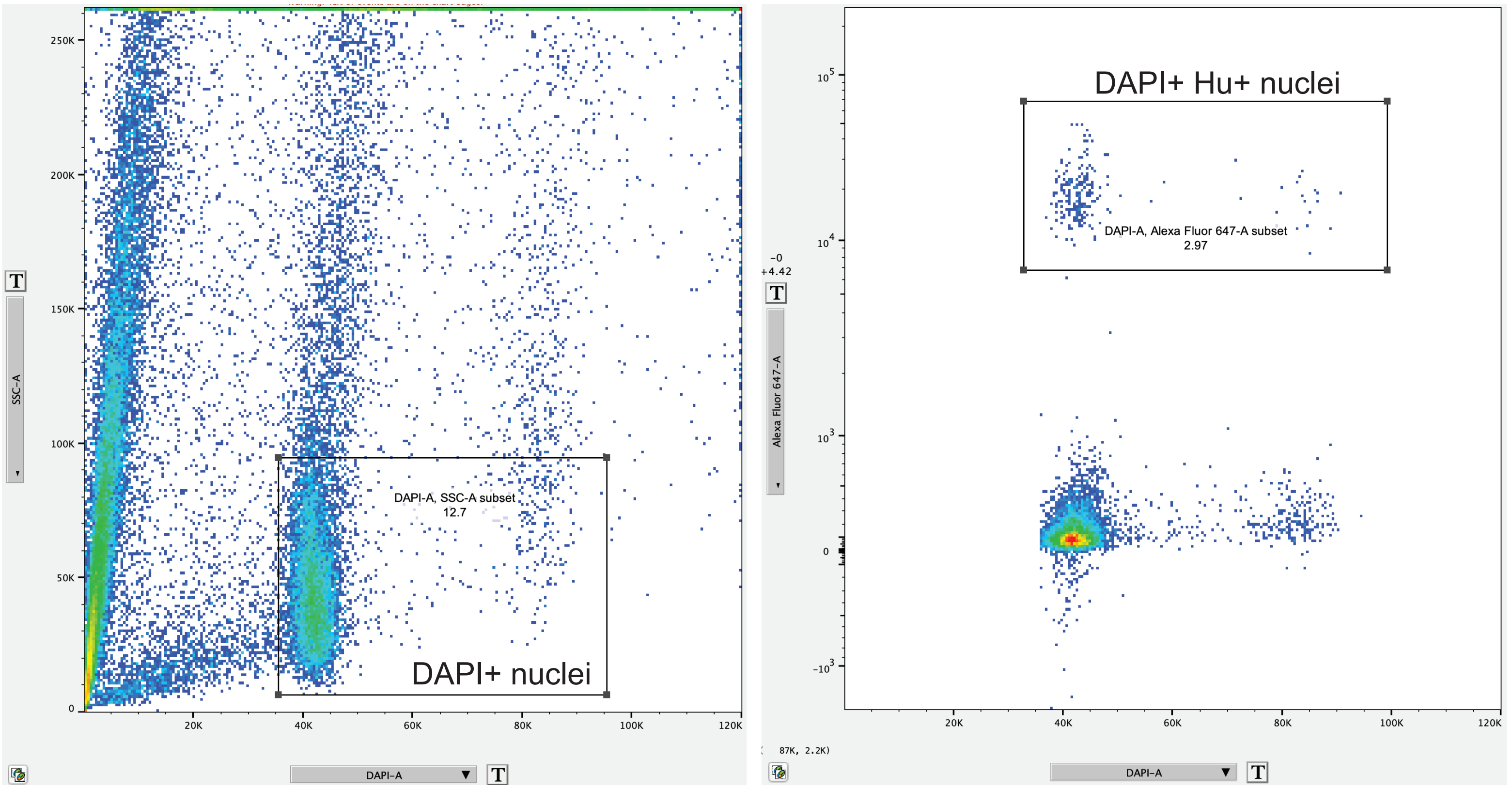
